# Consensus clustering applied to multi-omic disease subtyping

**DOI:** 10.1101/2020.10.19.345389

**Authors:** Galadriel Brière, Élodie Darbo, Patricia Thébault, Raluca Uricaru

## Abstract

**Background:** Facing the diversity of omic data and the difficulty of selecting one result over all those produced by several methods, consensus strategies have the potential to reconcile multiple inputs and to produce robust results.

**Results:** Here, we introduce ClustOmics, a generic consensus clustering tool that we use in the context of cancer subtyping. ClustOmics relies on a non-relational graph database, which allows for the simultaneous integration of both multiple omic data and results from various clustering methods. This new tool conciliates input clusterings, regardless of their origin, their number, their size or their shape. ClustOmics implements an intuitive and flexible strategy, based upon the idea of *evidence accumulation clustering*. It computes co-occurrences of pairs of samples in input clusters and uses this score as a similarity measure to reorganise data into consensus clusters.

**Conclusion:** We applied ClustOmics to multi-omic disease subtyping on real TCGA cancer data from ten different cancer types. We showed that ClustOmics is robust to heterogeneous qualities of input partitions, smoothing and reconciling preliminary predictions into high quality consensus clusters, both from a computational and a biological point of view. The comparison to a state-of-the-art consensus based integration tool, COCA, further corroborated this statement. However, the main interest of ClustOmics is not to compete with other tools, but rather to make profit from their various predictions when no gold-standard metric is available to assess their significance.

**Availability:** ClustOmics’ source code, released under MIT licence, as well as the results obtained on TCGA cancer data are available on Github: https://github.com/galadrielbriere/ClustOmics.

## Background

Recent advances in biological data acquisition made it possible to measure a wide range of data. Polymorphism data, DNA methylation, RNA expression, Copy Number Variations and other “omic” data are now routinely observed and analysed. Each omic type has the potential to reveal different molecular mechanisms associated to a phenotype, and making use of all available omic data could decipher complex and multi-level molecular interactions. Though several integrative tools have been developed, all of them aiming to answer biological questions by using multiple available data sources, the issue of omic data integration is far from solved. Along with the issue of omic data heterogeneity and integration, scientists are also challenged with the diversity of strategies and methods available to answer a same biological question, each approach having its own perks and benefits.

The question of cancer subtyping is particularly representative of this kind of issues. Disease subtyping aims at detecting subgroups of patients (samples) showing similar characteristics, by performing a clustering analysis. Even in the single-omic context such analysis can be challenging and numerous clustering strategies have been implemented and/or tested to this end: hierarchical clustering strategies, density-, distribution- or centroid-based strategies, supervised and unsupervised strategies, etc. As a matter of fact, the selection of a clustering method as well as of the optimal parameters to use, is generally tricky. Moreover, the various biological mechanisms that are involved may vary from one patient to another: each tumor is different and has its own characteristics, both in the tumor cells themselves and in their interaction with their environment. As these mechanisms are not circumscribed to a single molecular level, the detection of groups of patients showing similar characteristics across different omics is a key issue to enable personalised medicine, which aims to offer patients a treatment adapted to the characteristics of their tumors.

This motivated the development of new computational methods implementing different strategies to analyse simultaneously several omic datasets (for detailed reviews see [1, 2]). According to the classification proposed in [1], the *early integration* strategy consists in concatenating omic datasets in a large matrix and applying a clustering method conceived for single-omic data [3, 4]. On the other hand, *late integration* approaches first cluster each omic dataset independently and fuse single-omic clusterings into one multi-omic clustering [5, 6]. Other approaches perform *intermediate integration*, fusing sample similarities across omics [7, 8, 9], using dimension reduction strategies [10, 11], or statistical modelling with Bayesian frameworks [12, 13, 14].

To tackle both issues mentioned above, *i.e.*, multi-omic and multi-strategy integration, one may want to apply a particular type of late integration strategy by taking multiple clustering results (using different data, methods and parameters) and fuse all of them into one *consensus clustering*. Such a consensus clustering should benefit from the complementary information carried by various omics data and capitalise upon each method’s strengths while fading their weaknesses.

Note that with respect to classical late integration strategies that start from the raw omic datasets (*e.g.*, PINS [6] uses perturbations of raw omic data to generate the most stable multi-omic clustering), consensus clustering methods rely solely on clustering results. This is an essential property, as it allows for any clustering algorithm and any clustering result to be used, regardless of the availability of the raw omic dataset and of its type.

A naive way to compute a consensus clustering would be to perform the *intersection* of the clustering results, *i.e.* by simply taking the associations on which all methods agree. However, the greater the number of clusterings to fuse, the smaller the intersection. Moreover, when clusterings show different numbers of clusters, the question of the intersection is not trivial. Therefore, the issue of consensus clustering requires further methodological developments.

To compute a consensus clustering from a set of input clusterings, roughly two main strategies exist: object co-occurrence based approaches and median partition based approaches [15]. In the former strategy, consensus clustering is computed from a matrix counting co-occurrences of objects in the same clusters [5]. The latter strategy focuses on finding a consensus clustering maximising the similarity with the input partitions. Both strategies raise several non-trivial questions. The choice of a clustering algorithm and its tuning is not straightforward when working with a cooccurrence matrix. On the other hand, for the median partition based approach, the choice of a similarity measure is determinant. Nevertheless, for consensus clustering in a multi-omic, multi-method context, comparing co-occurrences of objects is more pertinent than comparing similarities between partitions.

Here, we present ClustOmics, a new graph-based multi-method and multi-source consensus clustering strategy. ClustOmics can be used to fuse multiple input clustering results, obtained with existing clustering methods that were applied on diverse omic datasets, into one *consensus clustering*, whatever the number of input clusters, the number of individuals clustered, the omics and the methods used to generate the input clusterings.

The co-occurrence strategy implemented in ClustOmics (detailed in the *Methods* section) is based on *Evidence Accumulation Clustering* (EAC), first introduced by Fred A. and Jain A. [16]. The idea is to consider each partition as an independent evidence of data organisation and to combine them using a voting mechanism. Similarly to clustering methods that use a distance or a similarity measure to compare objects, EAC considers the co-occurrences of pairs of objects in the same cluster as a vote for their association. The underlying assumption is that objects belonging to a *natural* cluster are more likely to be partitioned in the same groups for different data partitions. Thus, one can use the counting of the objects’ co-occurrences in clusters as a pairwise similarity measure. We further refer to these co-occurrence counts as *number of supports*. This measure, summarising the results from the input clusterings, is a good indicator of the agreement between the partitions and allows to produce a new partitioning that can be qualified as consensual. Although computationally expensive, this strategy allows exploiting all clustering results, regardless of the number of clusters, their size and shape.

We designed ClustOmics as an exploratory tool to investigate clustering results with the objective to increase the robustness of predictions, taking advantage of accumulating evidences. In order to allow the user to tackle a specific question and to explore relationship patterns within input clusterings and generated consensus, we store the data in a non-relational graph based database implemented with the Neo4j graph platform [17]. The use of a graph native database facilitates the storage, the query and the visualisation of heterogeneous data, hence allowing to develop a solution that is flexible to various integration strategies. Indeed, by fusing clusterings from different clustering methods, different data types, different experimental conditions, or several options at the same time, through the use of what we call *integration scenarios*, ClustOmics can address a wide range of biological questions. ClustOmics was applied in the context of multi-omic cancer subtyping, with TCGA data from different cancer types and multiple omic datasets. Input clusterings were computed with several single and multi-omic clustering methods and then were fused in a consensus clustering. Further details on the strategy implemented in ClustOmics are given in section *Methods*. In order to assess the benefit of this novel method, we further explored the robustness of our consensus clusterings with respect to the input clusterings, as well as their biological relevance, based on clinical and survival metadata available for each patient. We compare ClustOmics results with those of COCA [5], a well-known co-occurrence based consensus clustering tool that has already been used to combine multiple omic datasets to reveal cancer subtypes [18].

## Results

Consensus clustering for disease subtyping in a multi-omic context can be implemented as an *a priori* solution making a consensus of omic-specific input clusterings, or by *a posteriori* computing a consensus from multi-omic input clusterings. In order to better understand the perks and benefits of fusing omic data in a way or another, ClustOmics was tested in these two contexts based on two so-called *integration scenarios*.

First, we used ClustOmics to fuse multi-omic clusterings computed with existing integrative methods. In this scenario (Multi-to-Multi, MtoM), the integration of omics is performed by various existing clustering tools and ClustOmics’ integration computes a consensus result of the different multi-omic clusterings produced. The second scenario (Single-to-Multi, StoM) involves both methods and omics integration, as only single-omic clusterings computed from various methods are fused into one consensus clustering. See Figure 1 for a visual representation of these two scenarios.

**Figure 1.**
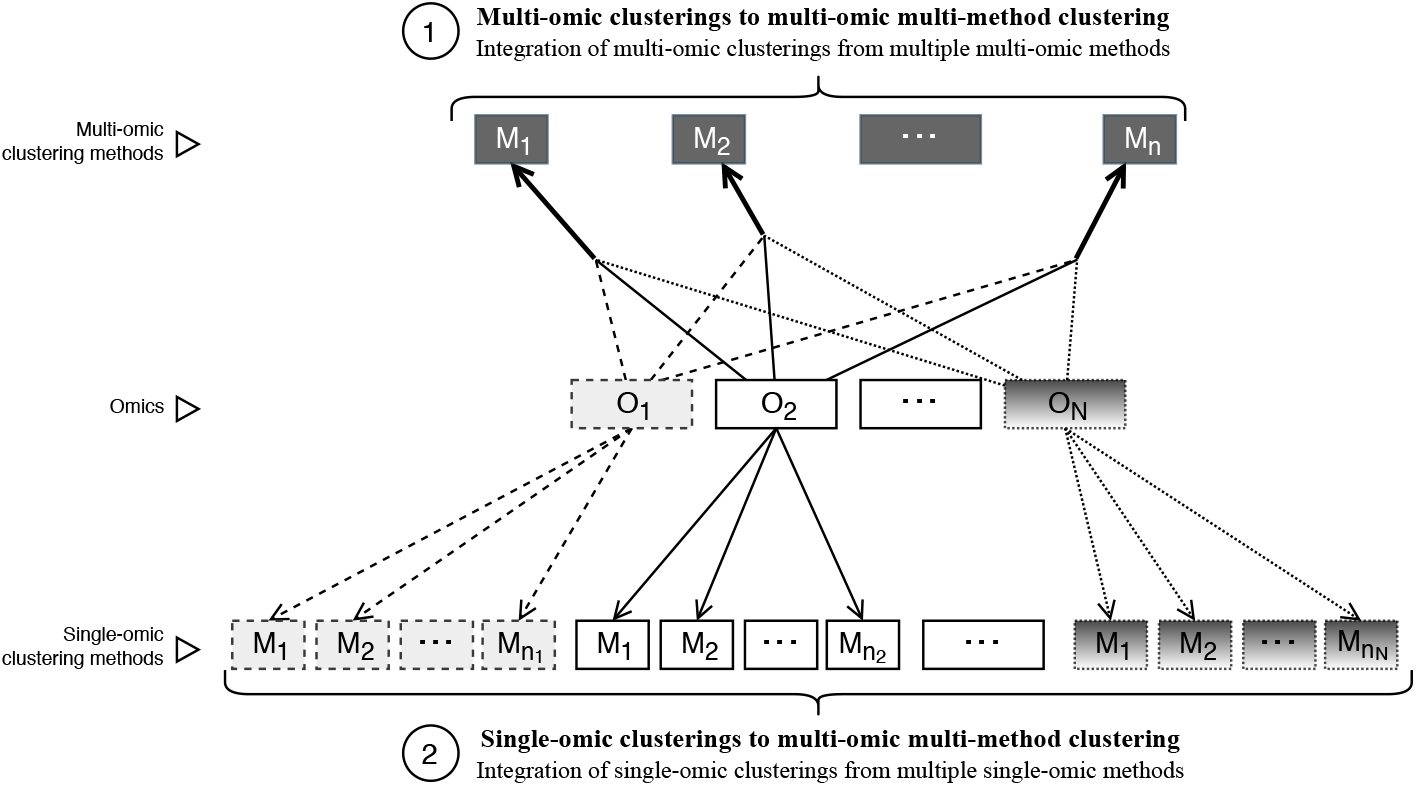
Two integration scenarios: Multi-to-Multi consensus clustering and Single-to-Multi consensus clustering. Arrows are dashed according to the omic considered by each input clustering method.

Below, we analyse and compare ClustOmics and COCA consensus multi-omic clusterings (produced using the same set of input clusterings) for the two integration scenarios, on TCGA data from ten different cancer types, three omic datatypes (gene expression, miRNA expression and methylation) and various input clustering strategies (described in section *Methods Datasets and tools used for computing input clusterings*). We further focus on breast cancer and analyse ClustOmics results for the Single-to-Multi scenario on the breast dataset.

### Results overview upon the ten cancer types for the two integration scenarios

Running ClustOmics and COCA on the ten cancer datasets with respect to the different integration scenarios implies starting by computing single- and multi-omic input clusterings to group patients according to their single- and multi-omic profiles. For the Multi-to-Multi scenario (MtoM), multi-omic input clusterings were obtained with existing multi-omic clustering tools: PINS [6], SNF [7], NEMO [8], rMKL [9] and MultiCCA [10] (see Table 6).

For the Single-to-Multi (StoM) scenario, the same tools listed above were applied, except for the MultiCCA tool that can only be used in a multi-omic context and that was replaced with the simple yet robust state-of-the-art method, k-means clustering [19]. In this scenario, the tools were applied on each omic dataset independently. Moreover, to evaluate the benefits of including patients with missing data (that were not measured for all of the three omics), two different runs were performed. In the first one, referred to as *StoM OnlyMulti*, only patients measured for the three omics were considered, that is patients with no missing data. For the second one, named *StoM All*, all available patients for each omic were kept. This implies that in this scenario the set of patients clustered in input clusterings was different across omics.

A survival and clinical label enrichment analysis was conducted on ClustOmics and COCA multi-omic consensus clusterings, as well as on the single-omic and multi-omic input clusterings (see section *Methods Biological metrics* for more details on the biological metrics used). An overview of the results upon the ten cancer types is displayed in Figure 2.

**Figure 2.**
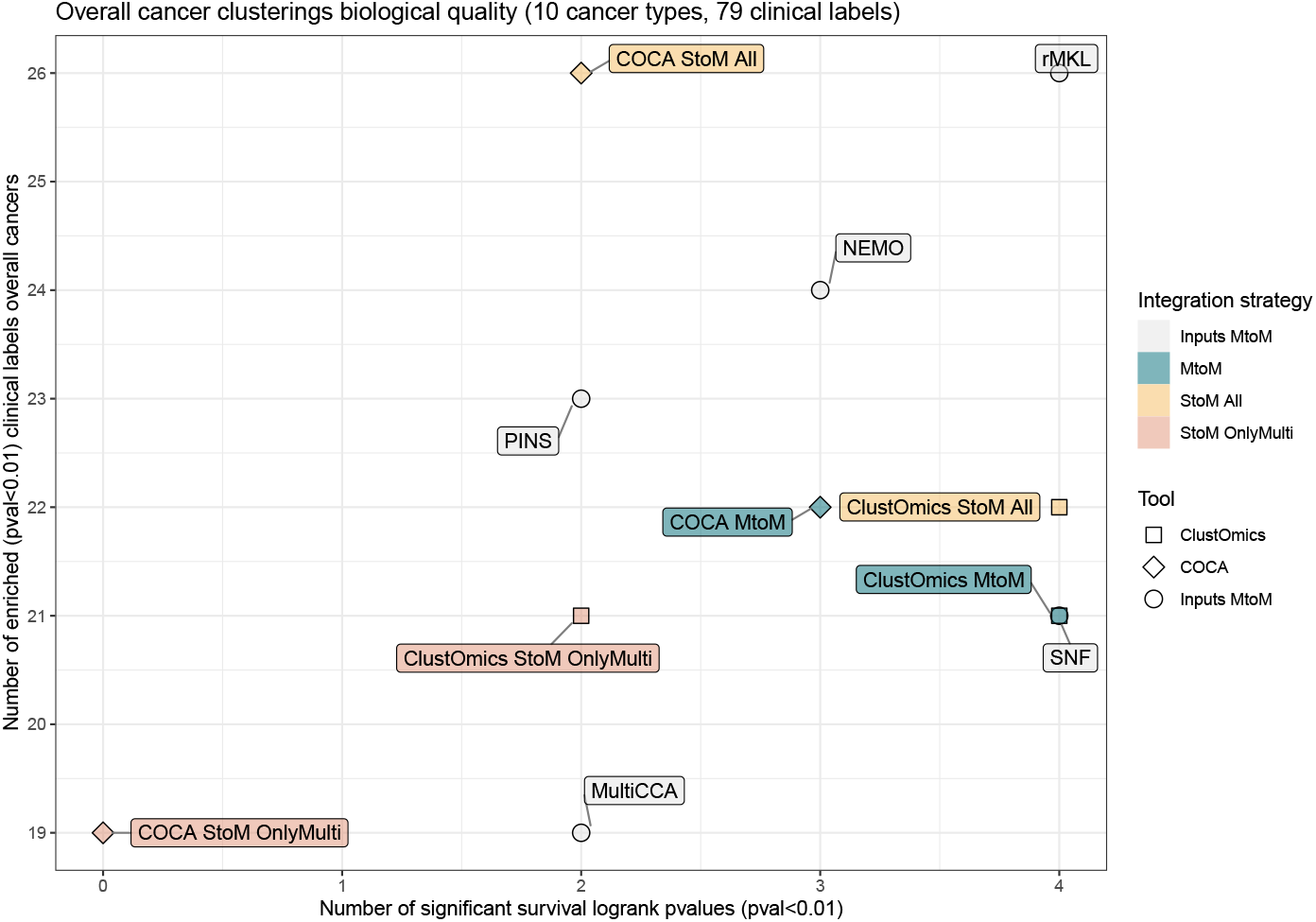
Overview of survival and clinical label enrichment results for the ten cancer types analysed. The *x* axis represents the number of significant survival p-values (< 0.01) found for each clustering, overall the ten cancer types. The *y* axis represents the total number of significantly enriched clinical labels (p-values < 0.01), all cancer types included. In total, 79 enrichment p-values were computed from 32 distinct clinical labels.

In terms of clinical label enrichment in clusters, the number of clinical labels significantly enriched varies from 19 to 26 (for a total of 79 enrichment p-values computed from 32 distinct clinical labels), depending on the clustering tool. The majority of clinical labels found enriched in ClustOmics consensus clusters were also found enriched for at least one input clustering as well as in the corresponding COCA consensus, and clinical labels stably enriched in input clusterings were also found enriched in ClustOmics and COCA consensus clusterings. For details with respect to the distribution of the clinical labels found enriched in input and consensus clusterings for the MtoM and StoM scenarios, see Supplementary Figure 1.

The survival analysis results show high heterogeneity. This supports the idea of computing a consensus clustering, especially when no gold-standard metric or ground-truth data are available. In this sense, it is important to stress out that ClustOmics succeeded to compute biologically relevant consensus partitions from input clusterings of variable quality. Indeed, in the Multi-to-Multi case, ClustOmics managed to find 4 out of 10 significant log-rank p-values, counterbalancing PINS and MultiCCA mitigated results, although they were part of the input clustering results used for this integration scenario. For the same set of input clusterings, COCA MtoM yielded to a 3 survival-wise significant consensus result.

Interestingly, for the Single-to-Multi scenario, Figure 2 clearly shows that considering individuals with missing data (StoM All) greatly improves the consensus clusterings, both in terms of clinical label enrichment and survival analysis, for ClustOmics as well as for COCA. For the StoM All scenario, COCA found 4 additional enriched clinical labels with respect to ClustOmics, but yielded to only 2 out of 10 survival-wise quality clusterings, compared to 4 for ClustOmics. Quality results for omic-specific input clusterings used for this scenario do not appear in Figure 2, but are further detailed in section *Results - Integration of single-omic clusterings*.

Detailed results for the MtoM and StoM integration scenarios are given in the following two sections.

### Integration of multi-omic clusterings (Multi-to-Multi scenario)

Input multi-omic clusterings were computed with the five multi-omic clustering methods presented in section *Methods - Datasets and tools used for computing input clusterings* using default parameters and following authors’ recommendations. The clusterings were produced using the multi-omic patients exclusively (those for which all three omic data are available). In order to make all input clusterings comparable, we run NEMO in the same way though, compared to the other five tools, NEMO is able to handle partial data.

ClustOmics was run with the *min_size_cluster* parameter arbitrarily set to 8 nodes for all cancer types, meaning that clusters of size below 8 were removed from the consensus clustering, the corresponding individuals being reassigned to consensus clusters exceeding the size threshold. We also set the *min_size_consensus* parameter to 95% of the population, to ensure that less than 5% of individuals are being reassigned to consensus clusters, either because of the number of supports threshold on the integration graph or because of the filter on the size of the clusters. Quantitative global measures on the ClustOmics consensus clusterings are detailed in Table 1.

**Table 1.** Scenario Multi-to-Multi - Number of patients initially clustered by ClustOmics, number of patients reassigned to consensus clusters, number of supports used to filter the integration graph, number of consensus clusters generated by ClustOmics and COCA, and average number of clusters in input clusterings.

| Cancer | # patients clustered | # patients reassigned | # supports threshold | # clusters ClustOmics | # clusters COCA | Avg # clusters inputs |
| --- | --- | --- | --- | --- | --- | --- |
| AML | 165 | 5 | 4 | 6 | 3 | 4.6 |
| BIC | 619 | 2 | 4 | 5 | 2 | 4.0 |
| COAD | 220 | 0 | 2 | 3 | 3 | 5.2 |
| GBM | 267 | 7 | 4 | 3 | 3 | 3.4 |
| KIRC | 176 | 7 | 3 | 3 | 3 | 4.2 |
| LIHC | 363 | 4 | 4 | 5 | 2 | 3.2 |
| LUSC | 341 | 0 | 2 | 2 | 2 | 3.4 |
| OV | 285 | 2 | 4 | 5 | 2 | 3.4 |
| SARC | 257 | 0 | 3 | 3 | 3 | 3.4 |
| SKCM | 351 | 0 | 3 | 5 | 3 | 4.8 |

Note that the maximum number of supports promoting the association of two patients in a same consensus cluster is bounded to 5, as five input clusterings (computed from five integrative clustering tools) were used for this integration scenario. After testing all possible thresholds on the number of supports, the optimal filtering threshold was obtained for each cancer type (2, 3 or 4, depending on the cancer type), meaning that only pairs of patients clustered in the same multi-omic cluster by at least 2 to 4 clustering methods were taken into account to compute the consensus clustering.

When comparing input and ClustOmics consensus clusterings, we observe a certain consistency in terms of the number of clusters. COCA on the other hand, resulted in 2 to 3 clusters independently from the cancer type, which suggests a lower sensitivity to input clustering dissimilarities compared to ClustOmics.

Two cancer datasets, COAD and LUSC, showed the lower consistency between the input predictions and clustered with a number of supports of 2. For LUSC cancer type, the consensus clustering resulted in only 2 clusters, despite the large size of the available cohort (341 individuals). The computation of the Adjusted Rand Index (ARI) [20] between input clusterings, a measure of similarity between partitions, showed that for these two cancer types, SNF and NEMO clusterings were very similar (with an ARI index of 0.7 for COAD SNF and NEMO clusterings and of 0.9 for LUSC, see Supplementary Figure 2) while the 3 other input clusterings showed high pairwise dissimilarity (*ARI* ≤ 0.4). Resulting consensus clusterings for both ClustOmics and COCA were very similar with SNF and NEMO and dissimilar with the other input clusterings, failing to compute an actual consensus of all input partitions. For the other cancer types, similarities between input clusterings were more balanced, enabling ClustOmics to reconcile predictions. ARI heatmaps comparing input and consensus clustering similarities for the ten cancer types in the MtoM scenario are available in Supplementary Figure 2.

Unsurprisingly, from Table 1 we remark that filtering the integration graph with a higher number of supports generally results in reassigning individuals (from 2 to 7) to consensus clusters. The predictions regarding these reassigned patients do not necessarily meet the number of supports threshold for which the consensus clusters were computed.

Figure 3 presents survival analysis results for the various multi-omic clusterings given as input to ClustOmics and COCA MtoM, and for the resulting consensus clusterings. When looking at the input clustering survival results, we can differentiate two cases:

**Figure 3.**
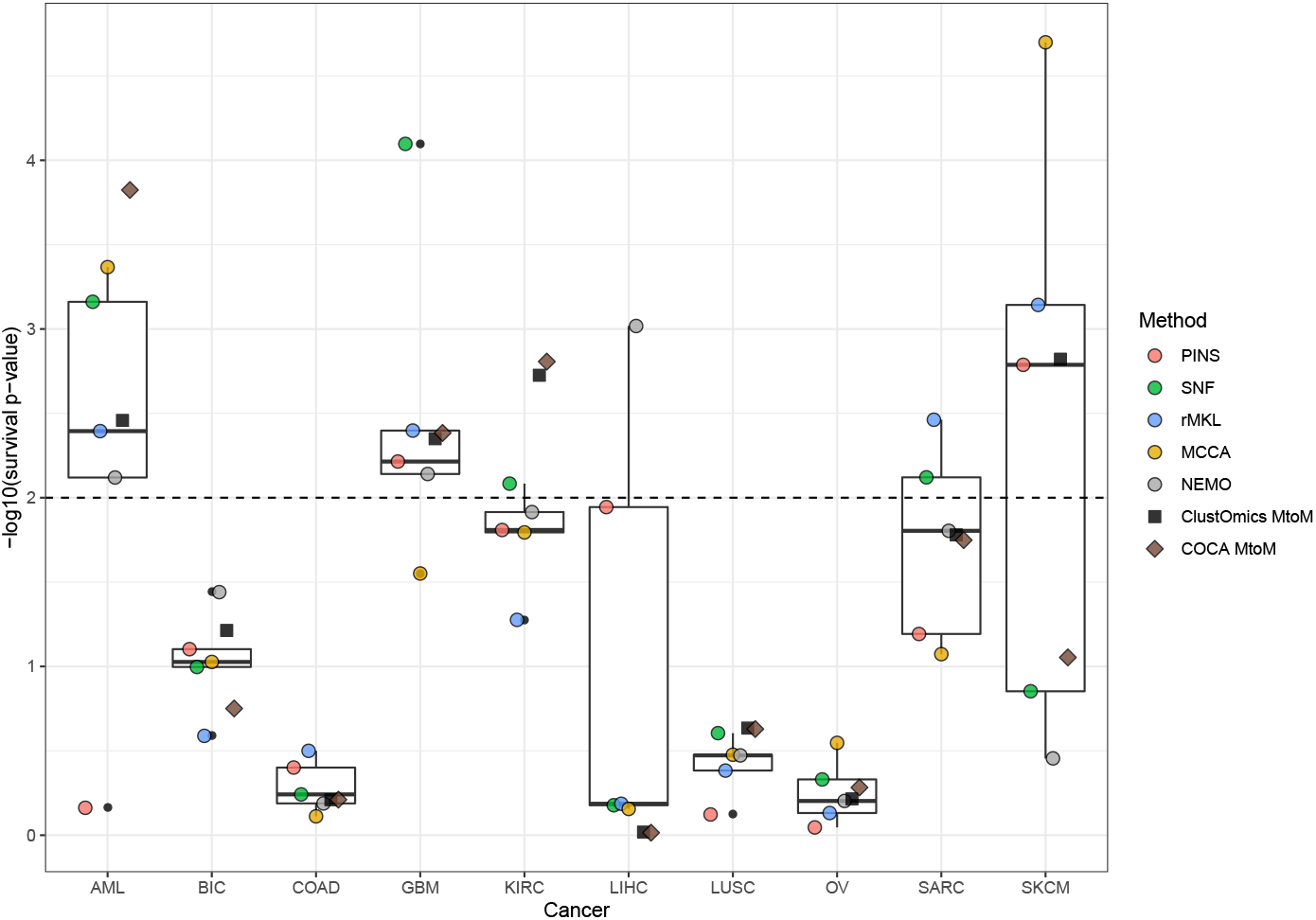
Survival analysis results for ClustOmics and COCA Multi-to-Multi consensus clustering and for each input multi-omic clustering. The horizontal dashed line indicates the threshold for significantly different survival rate (p-value ≤ 0.01). Boxplots were computed considering input clusterings only.

- For AML, LIHC, SARC and SKCM, input clusterings show a quite high heterogeneity in terms of survival quality
- For BIC, COAD, GBM, KIRC, LUSC and OV cancer types, input clusterings show relatively homogeneous survival quality

For the first group of cancer types, the heterogeneity of input clusterings survival qualities indicates how the choice of one clustering method can drastically impact the results. For these cancer types, ClustOmics produced consensus clusterings of a survival-wise quality approaching the median quality value, considering the input clusterings. Indeed, from input clusterings of various quality, ClustOmics was able to extract the most stable patterns across the input partitions.

When input partitions show homogeneous survival quality, ClustOmics gives similar results, which is an expected behaviour. The largest deviation from the median is found for KIRC cancer type, for which both COCA and ClustOmics consensus clusterings produced a partition of higher quality that could have been expected. Consensus clusterings were also investigated for clinical labels enriched in clusters. Upon the ten cancer types, ClustOmics and COCA found 20 common clinical labels as being enriched, of which 19 were also found enriched in at least one input clustering. On the other hand, 16 labels were found as enriched in at least one input clustering but not in the consensus clusterings (see Supplementary Figure 1-B). Table 2 give complete details on the clinical labels enriched in ClustOmics consensus clusterings for the ten cancer types.

For AML for example, ClustOmics computed clusters enriched for the CALGB cytogenetics risk category, a risk classification based on the Cancer and Leukemia Group B clinical trial [21], and for the French-American-British (FAB) morphology code, a clinical classification for AML tumors [22]. Reassuringly, BIC consensus clustering was found enriched for the PAM50 classification, a widely used breast-cancer subtype predictor [23].

**Table 2.** Clinical labels found enriched in Multi-to-Multi (MtoM) scenario consensus clusters, in Single-to-Multi (StoM All) scenario consensus clusters, and for both scenarios.

| Cancer | Scenarios | Enriched clinical labels |
| --- | --- | --- |
| AML | Both | Age at initial pathologic diagnosis<br>CALGB cytogenetics risk category<br>Leukemia French-American-British Morphology Code |
| BIC | Both<br>StoM All | Age at initial pathologic diagnosis, PAM50 call<br>Pathologic N, Pathologic Stage, Histological type<br>Estrogen receptor status, Progesterone receptor status<br>Pathologic M, Pathologic T |
| COAD | MtoM<br>StoM All | Histological type<br>Age at initial pathologic diagnosis |
| GBM | Both | None |
| KIRC | Both<br>StoM All | Pathologic M, Neoplasm histologic grade<br>Pathologic T |
| LIHC | Both | Gender, Age at initial pathologic diagnosis, Fetoprotein outcome value |
| LUSC | Both | None |
| OV | Both | None |
| SARC | Both<br>MtoM | Gender, Age at initial pathologic diagnosis, Histological type<br>New neoplasm event type |
| SKCM | MtoM | Age at initial pathologic diagnosis |

### Integration of single-omic clusterings (Single-to-Multi scenario)

To assess ClustOmics performance when fusing simultaneously input clusterings computed from different omic data and with different clustering methods, we investigated a second integration scenario, combining single-omic clusterings produced independently on each omic dataset. Overall cancer consensus results for this scenario are displayed in Figure 2, but discussed in detail in this section.

As stated above, single-omic clusterings were computed using the following five clustering tools: PINS [6], SNF [7], NEMO [8], rMKL [9] and k-means clustering [19] (with an optimal number of clusters computed with the Silhouette index [24]).

To assess the benefit of including individuals with missing data (not measured for all omics), two analysis were performed for this scenario:

- For StoM OnlyMulti, input clusterings were computed using exclusively multiomic patients. Given that this is the same set of individuals as in the MtoM integration scenario, the same parameters were used for all cancer types, *i.e.*, *min_size_consensus* = 95% and *min_size_cluster* = 8.
- For StoM All, omic clusterings were computed using all available patients. As the proportion of missing data varies between cancer types (up to 66% partial data for KIRC, see Table 5) and between omics, input clusterings do not apply to the same set of patients as in the scenarios previously described. In order to account for this increase in the number of patients to be clustered, *min_size_cluster* was set to 5% of the *multi-omic* population. *The min_size_consensus* parameter was set to 95% of the *multi-omic* population.

As shown in Figure 2, fully exploiting the available data (including patients with missing data) greatly improved the consensus clusterings, for both ClustOmics and COCA. Moreover, for 3 cancer types, BIC, GBM and LUSC, capitalising on all available individuals resulted in increasing the number of supports used to filter the integration graph. The largest increase in the number of supports threshold was observed for BIC, *i.e.*, from 7 supports in the StoM OnlyMulti up to 11 in the StoM All run. For LIHC, OV and SKCM on the other hand, we observe a decrease of, respectively, −2, −1 and −1 in the number of supports. For the other cancer types, the threshold on the number of supports is identical between the two runs. In the following of this study, we will focus on the results of the StoM All run.

In this scenario, the maximum possible number of supports is 15 as the five clustering methods were run on three omic datasets for each cancer. Note that for this scenario the threshold on the number of supports used to filter the integration graph has great influence on the capacity of ClustOmics to produce consensus clusters across omics and on the interpretation of the results. Indeed, the threshold has to be greater than 5 to ensure that all the conserved integration edges rely on an association that is consistent across at least two different omics (one omic being represented by five input clusterings). To ensure that all integration edges are built upon all three omics, the threshold must be 11 or higher. In fact, one should estimate an acceptable threshold depending on the experimental design and the biological question to address.

In our case, as we did not wish to bring any *a priori* on which omic should have a stronger impact on the results (indeed one omic data type could particularly well explain the disparities in molecular profiles of patients for a cancer type, but not for the others), we considered a number of supports of 7 to be sufficient to ensure that selected integration edges are either moderately consistent across the three omics, or strongly consistent in one omic. Together with the constraint to preserve at least 95% of the multi-omic population (*min_size_consensus* parameter), this gave a number of supports used to filter the integration graph ranging from 7 to 11. Results for this scenario are displayed in Table 3.

One of the major benefits of this integration scenario (besides the fact that single-omic clusterings are easier to compute) is its ability to cluster individuals that did not appear in all input partitions. Interestingly, although multi-omic patients have better chances to show high numbers of supports (as they appear in all input clusterings), some proportion of those multi-omic patients had to be reassigned to consensus clusters, while other individuals that were not measured for the 3 omics were clustered right away, which suggests a good agreement between the input clusterings for the classification of these individuals.

**Table 3.** Scenario Single-to-Multi All – Total population size (of which are multi-omic), number of patients clustered or reassigned to consensus clusters (of which are multi-omic), number of supports used to filter the graph, number of clusters generated by ClustOmics, number of clusters generated by COCA, and average number of clusters in the input clusterings.

| Cancer | Total<br>(multi-omic) | Clustered<br>(multi-omic) | Reassigned<br>(multi-omic) | # supports<br>threshold | # clusters<br>ClustOmics | # clusters<br>COCA | Avg #<br>clusters<br>inputs |
| --- | --- | --- | --- | --- | --- | --- | --- |
| AML | 197 (170) | 176 (163) | 21 (7) | 8 | 7 | 4 | 6,60 |
| BIC | 1096 (621) | 600 (600) | 496 (21) | 11 | 6 | 2 | 3,87 |
| COAD | 303 (220) | 276 (220) | 27 (0) | 7 | 6 | 4 | 3,47 |
| GBM | 578 (274) | 434 (262) | 144 (12) | 9 | 11 | 2 | 4,13 |
| KIRC | 534 (183) | 316 (174) | 218 (9) | 9 | 9 | 2 | 4,07 |
| LIHC | 377 (367) | 363 (354) | 14 (13) | 8 | 5 | 2 | 4,73 |
| LUSC | 501 (341) | 337 (321) | 164 (20) | 8 | 4 | 2 | 4,73 |
| OV | 591 (287) | 393 (272) | 198 (15) | 9 | 9 | 2 | 3,47 |
| SARC | 261 (257) | 261 (257) | 0 (0) | 8 | 3 | 3 | 4,33 |
| SKCM | 368 (351) | 345 (329) | 23 (22) | 8 | 6 | 4 | 4,67 |

Input clusterings can show great similarity for a given omic. If this omic allows to differentiate groups of individuals in a clear-cut way, it will drive consensus clustering. However, if the omic is less relevant to partition patients, input clusterings are more likely to show different patient associations. Such clusterings add noise-like integration edges in the integration graph, with low number of supports on edges. Therefore, we expect each omic to have a different impact on the final consensus clustering. To evaluate the impact of omic-specific input clusterings on the consensus result, we used the Adjusted Rand Index (ARI) [20].

In Figure 4, ClustOmics and COCA consensus clusterings were compared to each of the input clusterings. The relative proximity of a clustering consensus to the different input clustering, as measured by the ARI, indicates the tool’s ability to produce a partition that can genuinely be considered as reconciling the input predictions. In that respect, the ARI of a consensus clustering in relation to its inputs should be maximized for a maximum of input clusterings, including for those coming from different omics. The highest similarity between consensus and input clusterings from different omic sources is observed for the SARC cancer dataset (see Figure 4), with ARI values ranging from 0.4 up to 0.9 for at least one input clustering computed from each of the three omics datasets, suggesting similar associations at different molecular levels. For the other cancer types, the agreement between omic sources is less straightforward. Interestingly, for all cancer types, COCA and ClustOmics consensus clusterings resemble to the same input clusterings (computed from the same set of omic sources), thus suggesting that some omics are more appropriate to explain molecular differences between individuals. Unsurprisingly, gene expression impacts consensus clustering on most cancer types, but miRNA and methylation data also guided consensus clusterings, especially in COAD, LIHC and OV. Figure 4 also shows that the dispersion of ARI values is much greater for COCA consensus clusterings than for ClustOmics. While COCA consensus clusterings are very similar (if not identical) to few input clusterings but very dissimilar to the others, ClustOmics produces a consensus that is closer in average to all inputs.

**Figure 4.**
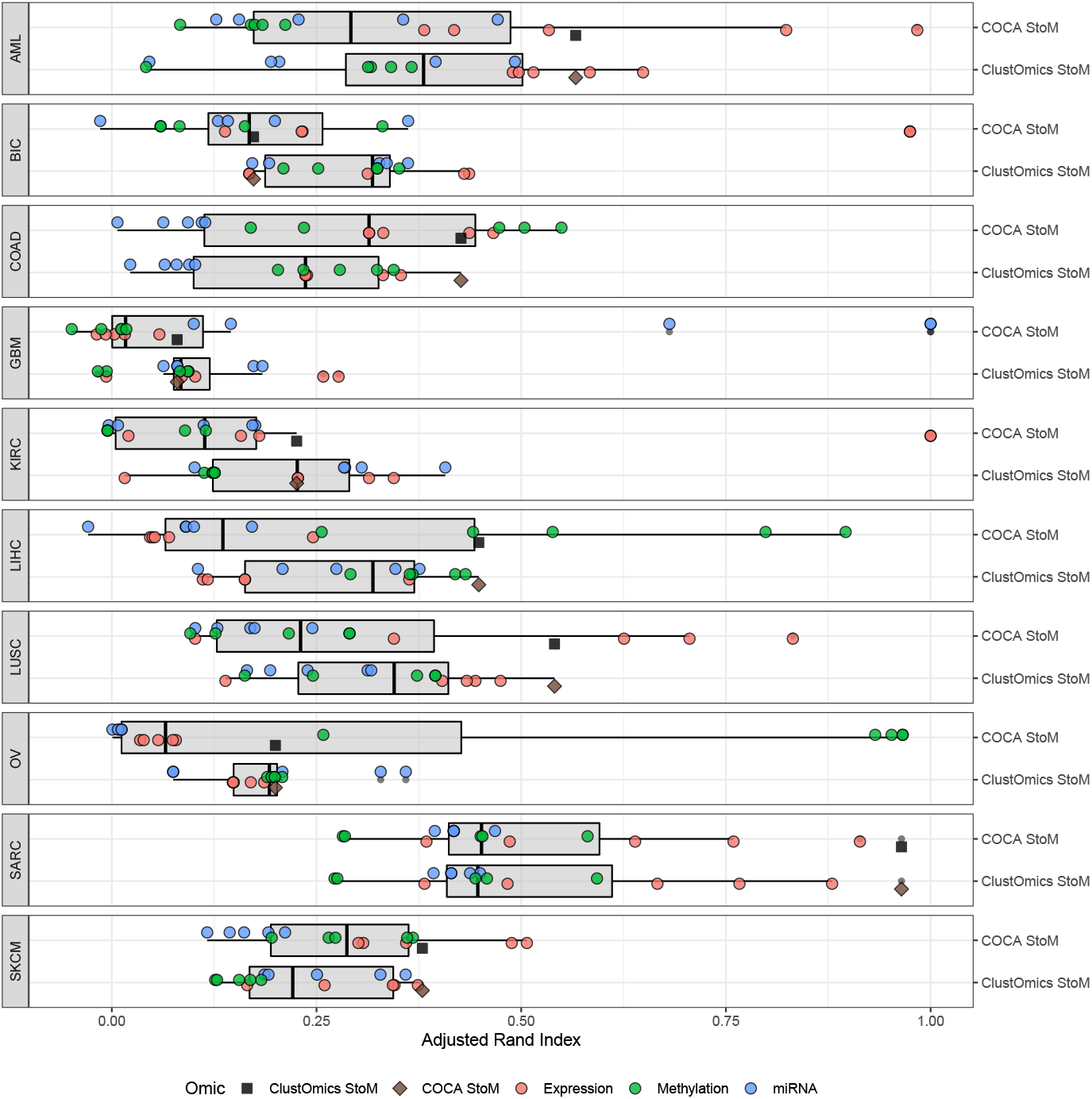
Adjusted Rand Index of input clusterings relative to ClustOmics and COCA StoM consensus multi-omic clusterings. Each point corresponds to one clustering, and is colored according to the omic type used. Each omic dataset was clustered using five different clustering tools (PINS, NEMO, SNF, rMKL, k-means) and therefore it is represented by five input clusterings. ClustOmics and COCA respective consensus clustering similarity is displayed with a black square and a brown diamond.

Survival analysis for this integration scenario (see Figure 5) shows two groups of cancer types, as already noticed for the Multi-to-Multi scenario. For BIC, COAD, LUSC and OV, gene expression, methylation and miRNA input clusterings show homogeneous survival p-values. For these cancer types, ClustOmics computed a consensus clustering with similar quality scores. For the cancer datasets showing higher heterogeneity among input clusterings survival quality, ClustOmics found significant survival p-values for AML, KIRC, LIHC and SKCM, despite some low quality clusterings that were given as input.

**Figure 5.**
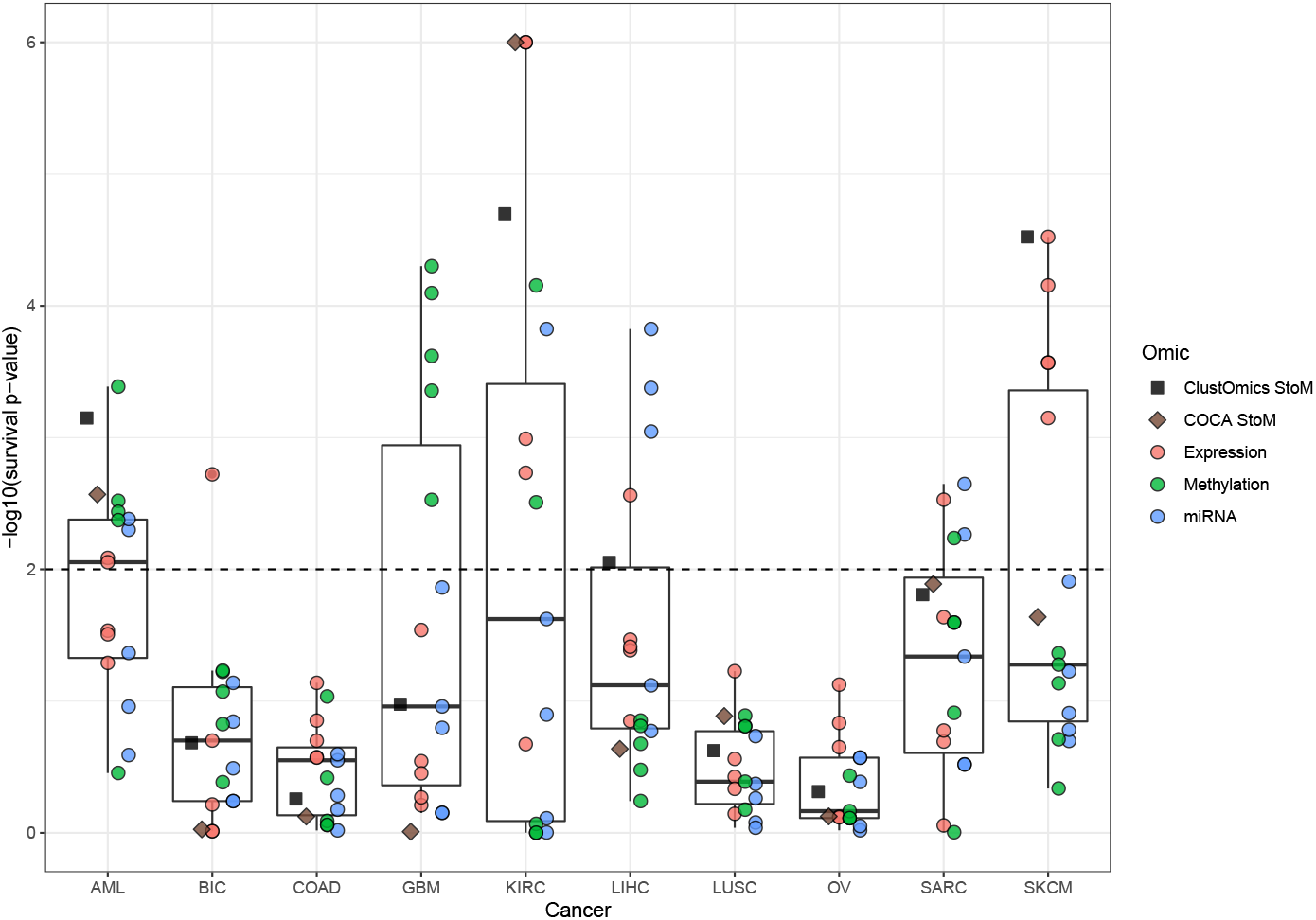
Survival analysis results for ClustOmics and COCA Single-to-Multi All consensus clustering and for each input multi-omic clustering. The horizontal dashed line indicates the threshold for significantly different survival rate (p-value ≤ 0.01). Boxplots were computed considering only input clusterings.

Clinical labels found enriched in consensus clusters are listed in Table 2. AML clusters were found enriched for both CALGB and FAB classifications, KIRC clusters for histologic grade and pathologic M and T (referring to the TNM classification of tumors [25]), LIHC clusters for gender, age at diagnosis and fetoprotein outcome value, while SKCM clusters showed no enriched clinical parameters. While BIC consensus clustering did not show good survival-wise results, pathologic M, N and T labels were found enriched in clusters, as well as pathologic stage, histological type, PAM50 call, and estrogen and progesterone receptor status.

In the following section we further explore the Single-to-Multi consensus clustering for BIC dataset.

### Study case: BIC Single-to-Multi consensus clustering

In this section we focus on the consensus clustering of the 15 single-omic clusterings for the BIC dataset (five clustering methods, listed in the previous section, applied on three omic data types) and analyse these results in parallel to the PAM50 classification. As the PAM50 classification is computed from the expression of 50 specific genes, while in this work we capitalise on three different omics, a certain heterogeneity in the clusters when compared to the PAM50 prediction is expected. Moreover, this heterogeneity is to be further explored, as it could reveal subtypes that are not distinguishable when considering only PAM50 genes, but that are heterogeneous when integrating other data sources.

From the 1096 patients available in BIC dataset (of which only 621 patients are measured for the three omics), ClustOmics succeeded to primarily classify 600 multi-omic patients in consensus clusters, with a number of supports threshold of 11 (see Table 3). The remaining 21 multi-omic individuals were reassigned to consensus clusters, as well as the 475 individuals with missing data. The consensus clustering resulted in a partition with 6 clusters, with sizes ranging from 115 to 254 individuals (the minimum allowed size for a cluster *min_size_cluster* being set to 5% of the multi-omic population, that is 31 individuals for BIC).

As the PAM50 clinical labels were missing for 255 patients, we applied the original classifier introduced by Parker *et al* [23] in order to call the missing labels. To estimate the quality of re-assessed PAM50 labels, we evaluated the concordance between available PAM50 labels and re-computed PAM50 labels. F1-scores showed Basal, Luminal A, Luminal B and Her2 PAM50 labels to be well predicted (F1-score of 0.89, 0.75, 0.74 and 0.64 respectively). Predictions for the Normal-like class are less reliable (F1-score of 0.27), due to the small size of the class (23 individuals). We further mapped the PAM50 calls to ClustOmics consensus clusters, and observed significant concordance, as depicted in Figure 6.

**Figure 6.**
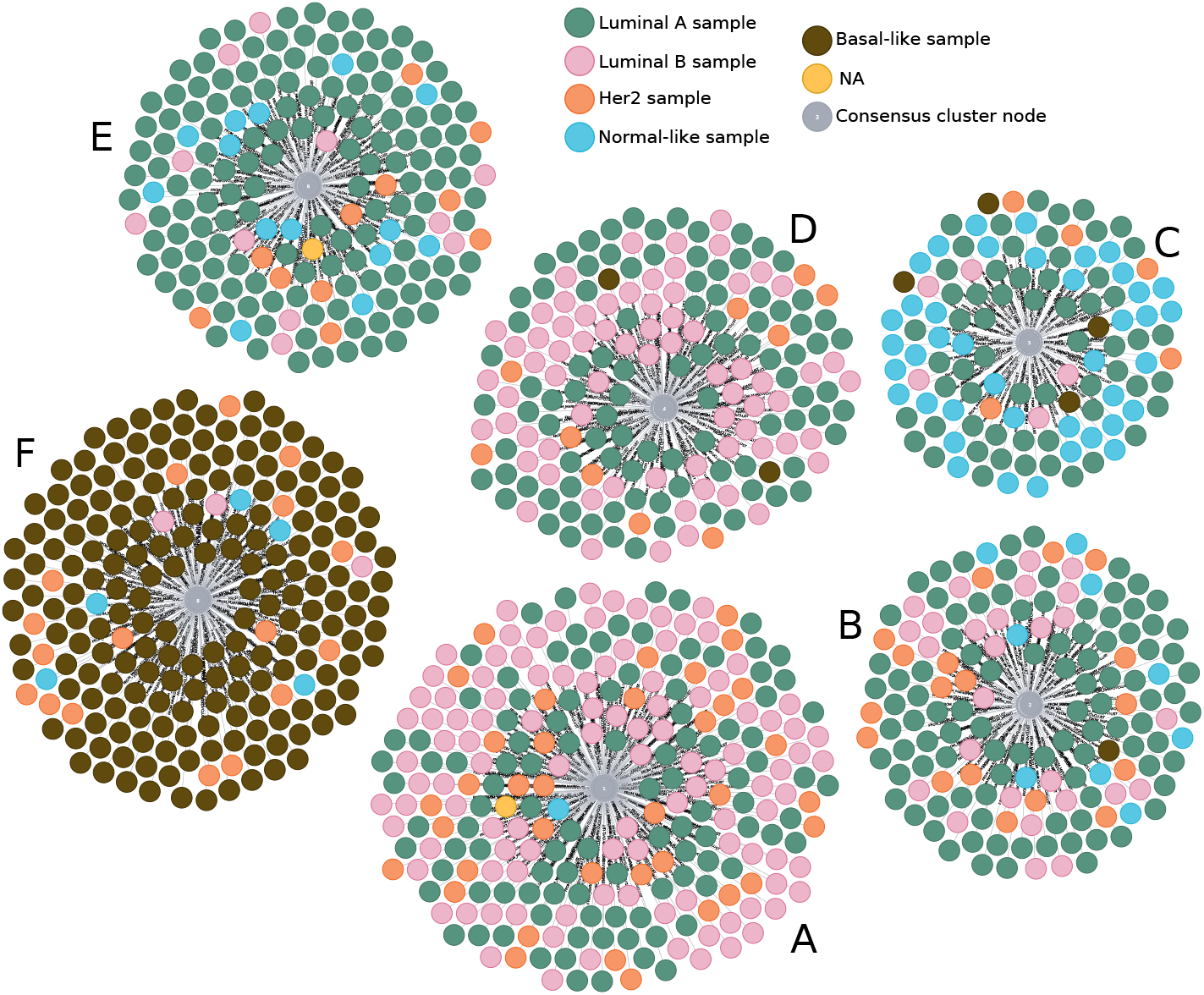
BIC consensus clustering with patients colored according to the PAM50 prediction. Annotated screenshot from Neo4j browser for graph visualisation.

Indeed, Luminal A samples are over-represented in the consensus clusters *B* and *E*, Luminal B samples in *A* and *D*, Her2 samples in *A*, and Normal-like samples in consensus cluster *C* (see Figure 6 and Table 4). The vast majority of Basal-like samples were classified in consensus cluster *F*, which gathers 190 of the 197 basal samples, the remaining 7 being clustered in consensus clusters *B*, *C* and *D*.

**Table 4.** Over and under represented clinical labels within BIC consensus clusters. ER+/ER− and PR+/PR− respectively correspond to Estrogen Receptor status and Progesterone Receptor status, positive and negative. M, N, T stages refer to the TNM staging system.

| Cluster | Over-represented labels | Under-represented labels |
| --- | --- | --- |
| A | Infiltrating Ductal Carcinoma<br>Her2, Luminal B<br>ER+ | Infiltrating Lobular Carcinoma<br>Basal, Normal-like<br>ER- |
| B | MX, N3, T3<br>Stage III<br>Infiltrating Lobular Carcinoma<br>Luminal A<br>ER+, PR+ | M0<br>Infiltrating Ductal Carcinoma<br>Basal<br>ER-, PR- |
| C | MX, T3<br>Infiltrating Lobular Carcinoma<br>Normal-like<br>ER+, PR+ | M0, T2<br>Infiltrating Ductal Carcinoma<br>Basal, Luminal B<br>ER-, PR- |
| D | Stage X<br>Mucinous Carcinoma<br>Luminal B<br>ER+, PR+ | Infiltrating Lobular Carcinoma<br>Basal, Normal-like<br>ER-, PR- |
| E | T1<br>Stage I<br>Luminal A<br>ER+, PR+ | Basal, Luminal B<br>ER-, PR- |
| F | N0<br>Stage II<br>Infiltrating Ductal Carcinoma, Medullary<br>Carcinoma, Metaplastic Carcinoma<br>Basal<br>ER-, PR- | Stage III<br>Infiltrating Lobular Carcinoma<br>Luminal A, Luminal B<br>ER+, PR+ |

**Table 5.** Number of patients measured per omic for each cancer type. Total: Number of patients measured for at least one omic (of which those having been measured for the three omics); proportion of partial data; Exp+Met: Patients measured for expression and methylation only; Exp+miRNA: Patients measured for expression and miRNA only; Met+miRNA: Patients measured for methylation and miRNA only; Exp: Patients measured for expression only; Met: Patients measured for methylation only; miRNA: Patients measured for miRNA only.

| Cancer | Total<br>(multi-omic) | % partial<br>data | Exp+<br>Met | Exp+<br>miRNA | Met+<br>miRNA | Exp | Met | miRNA |
| --- | --- | --- | --- | --- | --- | --- | --- | --- |
| AML | 197 (170) | 13,71 | 0 | 3 | 15 | 0 | 9 | 0 |
| BIC | 1096 (621) | 43,34 | 159 | 132 | 2 | 181 | 1 | 0 |
| COAD | 303 (220) | 27,39 | 57 | 0 | 0 | 8 | 18 | 0 |
| GBM | 578 (274) | 52,60 | 4 | 245 | 3 | 5 | 4 | 43 |
| KIRC | 534 (183) | 65,73 | 135 | 71 | 0 | 144 | 1 | 0 |
| LIHC | 377 (367) | 2,65 | 4 | 0 | 5 | 0 | 1 | 0 |
| LUSC | 501 (341) | 31,94 | 29 | 1 | 0 | 130 | 0 | 0 |
| OV | 591 (287) | 51,44 | 7 | 0 | 166 | 9 | 122 | 0 |
| SARC | 261 (257) | 1,53 | 2 | 0 | 2 | 0 | 0 | 0 |
| SKCM | 368 (351) | 4,62 | 16 | 1 | 0 | 0 | 0 | 0 |

**Table 6.** Methods used to compute input clusterings.

| Software | Multi-omic context | Single-omic context |
| --- | --- | --- |
| PINS [6] | Yes | Yes |
| SNF [7] | Yes | Yes |
| NEMO [8] | Yes | Yes |
| rMKL [9] | Yes | Yes |
| MultiCCA [10] | Yes | No |
| k-means [19] | No | Yes |

This mapping of PAM50 calls on consensus clusters, which seems fuzzy at first glance, is not surprising as it has been shown that separation of Luminal A and B samples was not reconstructed by RNA-seq unsupervised analysis [26]. Several studies also reported that the separation between Luminal subtypes was not consistent, suggesting that Luminal A and Luminal B samples may represent part of a continuum rather than distinct subgroups [26, 27, 28].

Moreover, clinical labels enrichment analysis and additional tests applied to describe the clusters show good mapping between ClustOmics clusters and key biological clinical labels, such as Estrogen and Progesterone Receptor status (ER/PR) or histological type of tumors (see Table 4).

Finally, we investigated the biological relevance of ClustOmics consensus clustering by comparing gene expression profiles between clusters. We computed the top 1000 genes differentially expressed across groups by applying the Kruskal-Wallis test [29] and selecting FDR adjusted p-values bellow 0.001 (see Supplementary Figure 3). We clustered the top 1000 genes in 6 clusters using hierarchical clustering and for each gene list, we looked for over-represented Biological Process (BP) related Gene Ontology terms (GO terms). One of the gene clusters showed no significant results (FDR adjusted p-values ≥ 0.05), but the other 5 gene lists were found enriched for cilium organisation and assembly, response to transforming growth factor *β*, tissue migration, T-cells activation, mitotic nuclear division or other biological processes (see Supplementary Figure 4).

## Discussion

The novel method we present in this paper deals with two key issues raised by the present context in biology and medicine and, in parallel, in bioinformatics. Indeed these domains witness an actual revolution in the acquisition of molecular data thus facing a flood of various types of omic data. The ultimate goal is to benefit from the diversity and complementarity of these omic data (DNA methylation, Copy Number Variations, polymorphism data, etc.), by analysing them simultaneously. However, multi-omics data integration is only a facet, as at the same time we face an outburst of biocomputational approaches meant to deal with this unprecedented variety and quantity of data, and the choice of a method or of the optimal parameters is generally challenging. In this paper both simultaneous integration of multiple omics and of various methods are tackled in an innovative manner through an original integration strategy based on consensus.

More particularly, in this work we address the cancer subtyping problem from a personalised medicine related perspective that is gaining increasing attention. In order to treat patients according to their disease profile, one should be able to distinguish between disease subtypes. These can be predicted from omic data (traditionally gene expression, but also methylation, miRNA, etc.) by performing patient clustering (hierarchical clustering, density-based clustering, distribution-based clustering, etc.). Our novel graph-based multi-integration method, can fuse multiple input clustering results (obtained with existing clustering methods on diverse omic datasets) into one consensus clustering, whatever the number of input clusters, the number of objects clustered, the omics and the methods used to generate the input clusterings.

To compute a consensus clustering, our method, implemented in a tool called ClustOmics, uses an intuitive strategy based on evidence accumulation. The evidence accumulation counts (*i.e.*, the number of supports on the integration edges) make consensus clustering results easier to interpret, as they add insight to the extent to which the consensus clustering can be considered as being multi-source (issued from multiple omics) and to the overall agreement of input partitions.

The original EAC strategy as proposed by Fred and Jain [16] uses input partitions obtained by running the k-means algorithm multiple times (≈ 200) with random initialisation of cluster centroids. From these partition results, a co-occurrence matrix is computed and a Minimum Spanning Tree algorithm is applied to find consensus clusters, by cutting weak links between objects at a threshold *t* defined by the user. The authors recommend that clusterings obtained for several values of *t* should be analysed. In ClustOmics, we developed a weighted modularization optimisation strategy to automatically select the best filtering threshold. Also, rather than generating input clusterings from running multiple times the same algorithm as proposed by Fred and Jain, here we make benefit from using various clustering strategies, each searching for different patterns and giving different insights to the data. This also allows the use of algorithms that are specialised for one omic. Moreover, by taking as input a high number of clusterings obtained with a same tool with varying parameters, the convergence of the consensus clustering, especially in a single-omic context, can be improved and this can also be achieved by ClustOmics by giving the appropriate input clusterings.

It should be noted that though our method does not formally weight input datasets (*e.g*, according to their level of confidence), one can artificially enhance the impact of one or several omic sources by providing supplementary single-omic input clusterings. In the same way, when dealing with missing data, patients measured in all omics are more likely to accumulate supports and therefore more likely to cluster together. In a context of multi-source integration, favoring individuals with the least amount of missing data makes it possible to highlight the predictions supported by several data sources, which is the desired behavior. In a context of single-source integration, the same set of objects is usually used in all input clusterings (apart from a few specificities of the input clustering tools used).

TCGA real datasets from three different omics and ten cancer types were analysed with respect to two integration scenarios: (1) fusing multi-omic clusterings obtained with existing integrative clustering tools and (2) fusing omic-specific input clusterings. In both cases, ClustOmics succeeded to compute high quality multi-omic consensus clusterings with clusters showing different survival curves and enriched for clinical labels of interest, coherent with what could be found in the cancer literature. Moreover, results indicate that ClustOmics is robust to heterogeneous input clustering qualities (reconciling and smoothing the disparities of partition) and in comparison with a state-of-the-art consensus based integration method, COCA. Notably, our method is not meant to compete with existing single or integrative omics clustering methods but, on the contrary, it aims at capitalising on the preliminary input predictions in order to increase their robustness by taking advantage of accumulating evidence to reveal sharper patterns in the data. This selection of robust patterns across input partitions renders ClustOmics stable when facing heterogeneous input clusterings, and particularly useful when no gold-standard metric is available to assess the quality of the results. Hence, with a sufficient number of input clusterings, no prior analysis of the input is needed, given that low quality clusterings, likely to add noise to the integration graph, play a smaller part in the evidence accumulation. Omics for which the separation of samples is clear-cut will drive the consensus clustering while omics that do not show interesting patterns across samples will be faded via the integration graph filtering step. For the same reason, it is important to highlight that, as long as the signals in the available omics are strong, ClustOmics is able to cluster samples that do not appear in all omic datasets, making use of available data and addressing the issue of partial data.

Finally, though presented in a disease subtyping context, one should grasp that our method is not limited to this application case. Indeed taking partition results (instead of raw data) as input, makes it generic and adaptable to a wide range of biological questions, as one can use any kind of partitioning of the data, *e.g.* clinical labels, groups of genes of interest, etc., as an input clustering. A major strength of ClustOmics resides in its exploratory aspect, resulting on one side from a flexible intrinsic model that gives the user complete power on the integration scenario to investigate and, on the other side, from the use of the graph-oriented database Neo4j. All input data and metadata are stored in this kind of database, which may be easily queried and visualised by a non-specialist with the Neo4j browser.

## Conclusion

Facing the diversity and heterogeneity of omic data and clustering strategies, one might want to make profit from all available data to compute a consensus clustering. ClustOmics is able to fuse any set of input clusterings into one robust consensus, which can easily be interpreted based on the number of supports evidence accumulation score. ClustOmics can be adapted to answer a wide range of biological questions. The use of integration scenarios allows users to explore various integration strategies, by adding or discarding data sources and/or clustering methods.

### Methods

In this section we detail the strategy implemented in ClustOmics. We then describe the datasets and the metrics that were used to evaluate our new method. For the sake of simplicity, as in this paper ClustOmics was applied in the context of cancer subtyping, we will further refer to objects of interest as *Patients*. However, ClustOmics can be applied on different biological entities, like genes or cells for instance.

### ClustOmics integration strategy

ClustOmics integration strategy, depicted in Figure 7, starts from a set of input clusterings generated with various clustering methods and/or from different omic sources.

**Figure 7.**
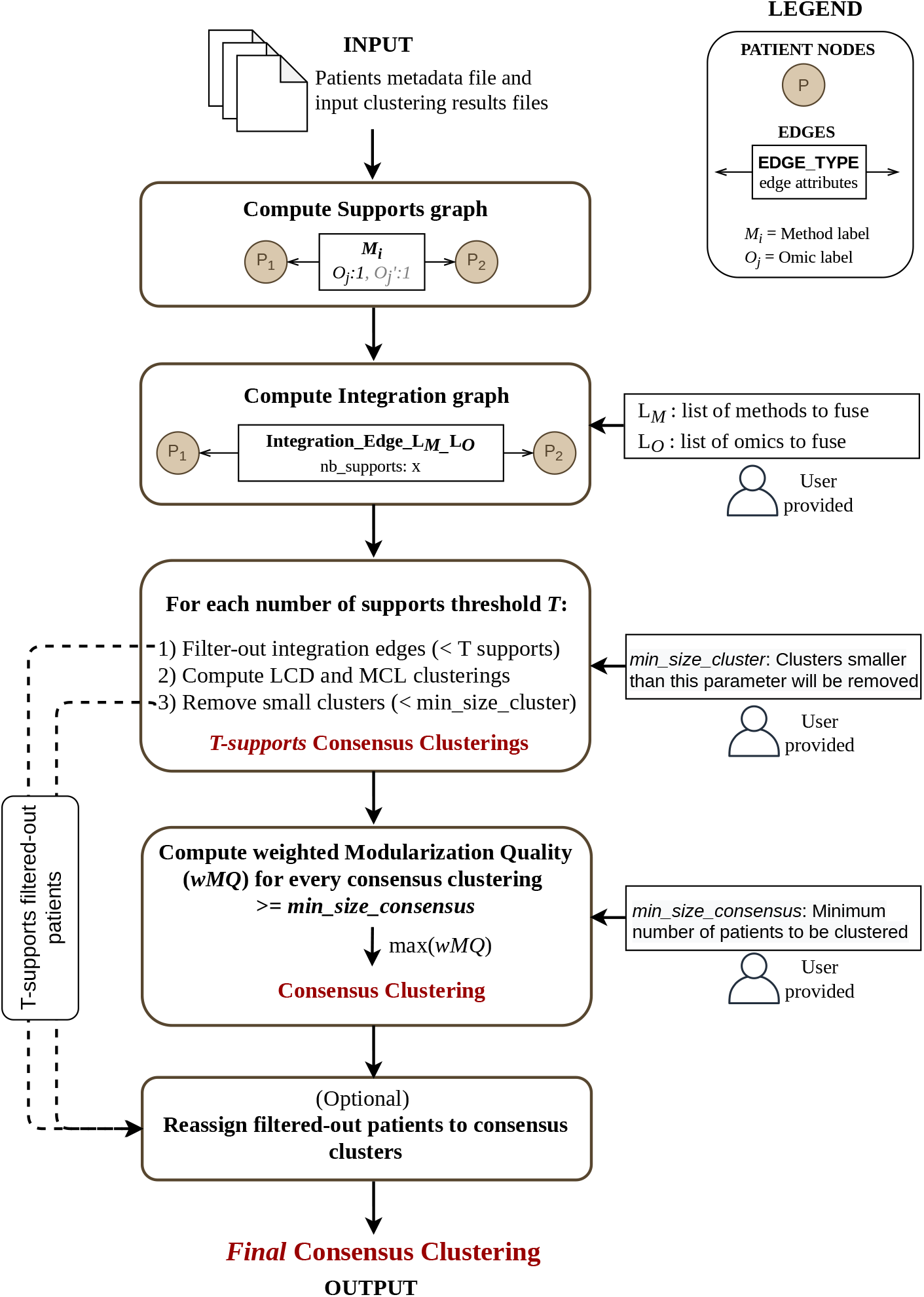
An overview of the strategy implemented in ClustOmics.

First, from a patients metadata file and from available input clusterings, a Supports graph (SG) is instantiated. In this graphs each *Patient (P)* corresponds to a node and shares a *Support edge* with another patient when classified in the same cluster (co-clustered) in at least one input clustering. One support edge relates to one input clustering tool, and each support edge displays one or multiple attributes to indicate the omic sources supporting the co-clustering of the patients.

Next, given an integration scenario, meaning a list of omics and methods to integrate, the corresponding *Integration Graph (IG)* is computed. Then, the integration graph is filtered and clustered to produce a consensus clustering according to the given integration scenario.

Below, we detail the integration graph computation, filtering and clustering steps, resulting in a ClustOmics consensus clustering.

### Compute the Integration Graph (IG)

Given an *Integration Scenario* (defined by a set of input clusterings), ClustOmics exploits the information on the support edges to compute the so-called *number of supports*, by counting the considered input clusterings sustaining the association of patients. The numbers of supports are reported on the *Integration Edges* and so, for a given integration scenario, a pair of patients may share at most one integration edge. In this way, heterogeneous data is aggregated into co-occurrence counts that are used as a similarity measure to perform Evidence Accumulation Clustering (EAC) [16].

### Filter and cluster the Integration Graph

Integration graphs are generally densely connected, as each pair of nodes may have been clustered together at least once over the set of omics and methods. However, as integration edges are weighted with the number of supports agreeing on the corresponding associations, the most robust integration edges can be distinguished from predictions that are not consistent across omics and methods. Hence, ClustOmics filters the graph according to the number of supports by removing non-consistent integration edges. The goal is to obtain a filtered graph foreshadowing *natural* clusters that correspond to a consensus. The choice of a threshold to filter the integration edges is therefore determinant.

Figure 8 depicts the impact of an increasing number of supports filtering threshold on the internal structure of the integration graph. One can observe that two issues arise from this filtering process:

- First, increasing the threshold generates smaller graphs. Indeed, pairs of nodes that do not share any integration edge with a sufficient number of supports are removed, leading to a partial classification of the input set of patients.
- The second issue is the loss of structure in the filtered integration graph: when filtering at a high threshold, the resulting graph may become too sparse to be considered informative, like the graph in Figure 8-c with numerous small connected components.

**Figure 8.**
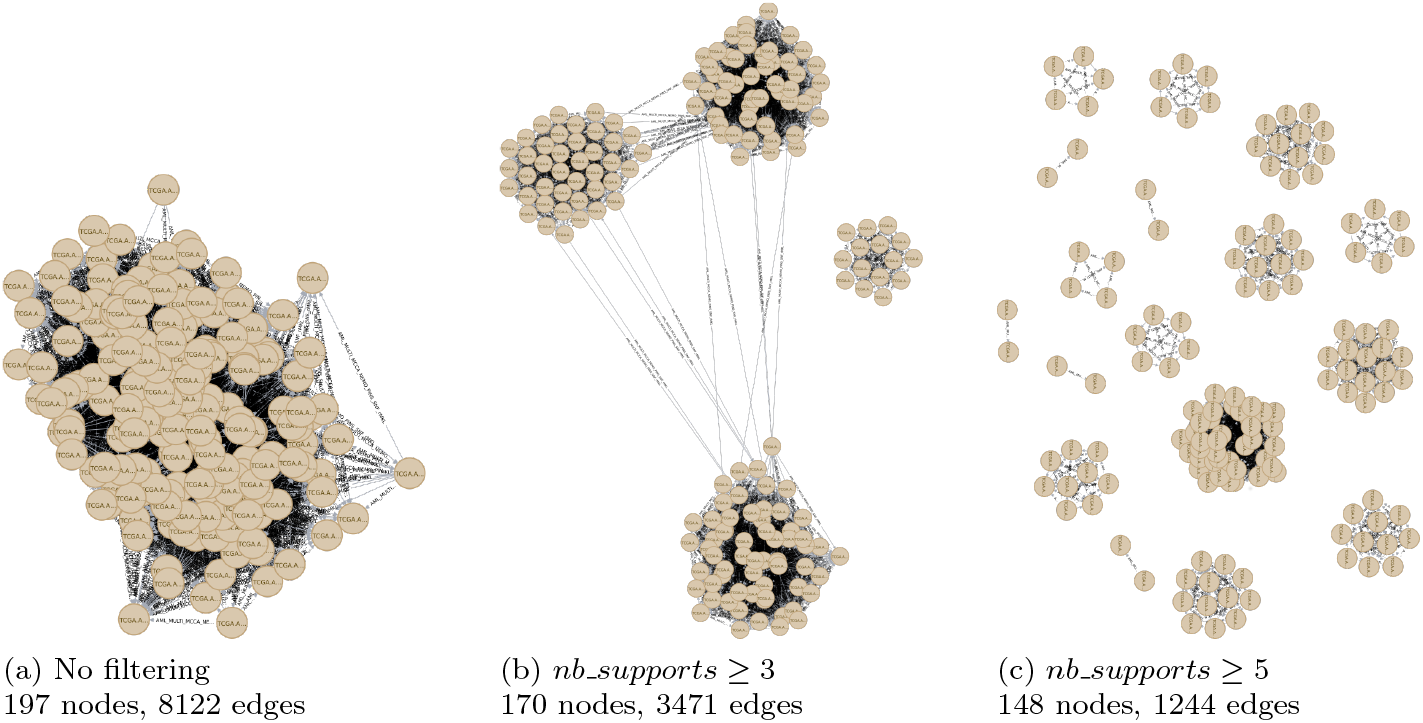
An integration graph filtered with increasing threshold values: 1, 3, 5 (the maximum number of supports for an integration edge being 5 in this example). Screenshots from Neo4j browser for graph visualisation.

Therefore, producing a relevant classification requires finding the best compromise for the support threshold. For this, ClustOmics tests all possible configurations by iteratively filtering the integration graph with increasing support thresholds. At each iteration, it uses state-of-the-art graph clustering methods (see the subsection below) to compute consensus clusterings for the corresponding filtered integration graph.

It is important to note that the resulting consensus clusterings should be analysed with respect to the number of supports used to filter the graph prior to clustering. This number of supports indicates the level of agreement between the input clusterings and gives insight on the extent to which the resulting consensus clustering can be considered as being truly multi-omic.

Moreover, in order to deal with the two issues described above and to keep merely informative results, each integration graph consensus clustering result goes through an additional filtering step with the two following parameters:

- the *min_size_cluster* parameter indicates the minimum accepted size for a cluster that is part of the consensus clustering. Clusters with less than *min_size_cluster* nodes are removed from the analysis.
- the *min_size_consensus* parameter indicates to what extent ClustOmics is allowed to discard nodes, *i.e.*, patients. ClustOmics will further consider only consensus clustering results having at least *min_size_consensus* nodes.

Finally, a quality metric, *i.e.*, the Weighted MQ index, is computed for all consensus clusterings that passed the filtering steps.

Optionally, one may want to reconsider the individuals that were discarded during the filtering steps (either when filtering-out integration edges or small clusters) and analyse them with respect to the consensus clusters. With this in mind, ClustOmics is able to reassign filtered-out individuals with respect to the mean number of supports shared with patients from consensus clusters, though such additional predictions do not necessarily meet the threshold with which the consensus clusters were originally computed.

Below we give insights on the graph clustering algorithms that are used to compute the consensus clusters, as well as on the quality metric employed for the identification of a robust consensus clustering, the *weighted Modularization Quality*.

### Graph clustering algorithms

ClustOmics filters the integration graph for each possible threshold on the number of supports and, for each filtered graph it computes two consensus clusterings with two state-of-the-art, complementary graph clustering algorithms: the Louvain Community Detection (LCD) algorithm, based on modularity optimisation [30], and the Markov Clustering (MCL) algorithm, based on the simulation of stochastic flow in graphs [31].

Modularity optimisation is one the most popular strategies in graph clustering algorithms [32], while MCL/MCL-based methods have proven highly efficient in various biological network analysis (protein-protein interaction networks [33, 34], protein complex identification [35], detection of protein families [36]). Moreover, it has been shown that modularity optimisation algorithms present a resolution issue [37, 38]: a tendency to fuse small clusters (even for those that are well defined and have few inter-connections) thus favoring the formation of bigger clusters than those computed by MCL [39]. Small clusters predicted by MCL can be an issue in ClustOmics case, as it removes clusters smaller than the user-defined *min_size_cluster* parameter, considering them to be non-informative.

### Selection of the best consensus clustering based on the Weighted MQ index

The *Modularization Quality* (*MQ*) was first defined by *Mancoridis et al.* in the context of software engineering [40]. Compared to the popular *Modularity* measure [41] optimised in graph clustering algorithms, which compares the distribution of edges with respect to a random graph with the same number of vertices and edges as the original graph, the *Modularization Quality* (*MQ*) evaluates the quality of a clustering as the difference between internal and external connectivity ratios. That is, the ratio between the number of connections observed within a given cluster and between two given clusters, and the maximum possible number of such edges. An optimal clustering for this measure should maximise the intra-connectivity ratio (every two nodes belonging to the same cluster share an edge) and minimise the inter-connectivity ratio (nodes classified in different clusters do not share edges). Indeed, in the context of consensus clustering based on evidence accumulation, it makes more sense to compare the distribution of the edges in the integration graph to the case where all nodes would have been partitioned in the same, optimal way in all input clusterings, *i.e.*, a graph where all intra-cluster nodes are connected and all inter-cluster nodes are disconnected.

Moreover, we adapted the original *MQ* index for weighted, undirected graphs with no self loops (in our case a *Patient* node cannot share an integration edge with itself). We denote this adaptation of the Modularization Quality, as the *weighted Modularization Quality* (*wMQ*).

Let *G* = (*V, E*) be a graph where *V* denotes the set of nodes and *E* the set of edges of *G*. Let *C* = (*C*_1_,…, *C_k_*) be a consensus clustering with *k* clusters and |*C_i_*| the number of nodes classified in cluster *C_i_*. Let us also note *w*(*e*) the weight of a given edge, *W* (*e_ii_*) the sum of weights of the edges internal to *C_i_* cluster (connecting vertices from *C_i_*), *W* (*e_ij_*) the sum of weights between clusters *C_i_* and *C_j_* (connecting a vertex from *C_i_* to a vertex from *C_j_*) and *max*(*w_IG_*) the maximum possible weight on the edges of the given integration graph (the maximum possible number of supports, also corresponding to the number of input clusterings being fused). We therefore define the *wMQ* index computed for a consensus clustering *C* obtained on the integration graph *IG* as:

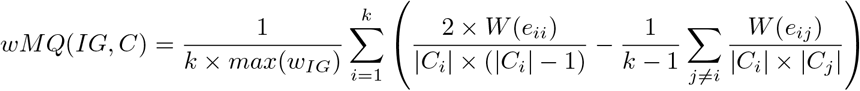

The first term of the sum corresponds to the weighted internal connectivity ratio for a cluster *C_i_*. Indeed, the sum of the internal edges weights *W* (*e_ii_*) is adjusted with the maximum possible value of the sum of the edges linking a set of |*C_i_*| nodes, which would be reached if all *C_i_* nodes were connected with *max*(*w_IG_*) weighted edges. Note that for an undirected and no self-loop graph, the maximum number of edges in a subgraph of |*C_i_*| nodes is 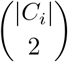. Similarly, the second term of the sum represents the weighted external connectivity ratio of a cluster *C_i_*, given by the sum of the weights of the edges linking a node from cluster *C_i_* to a node belonging to a cluster *C_j_* (≠*C_i_*).

The *wMQ* values range from −1 to 1, where a *wMQ* of −1 corresponds to the case where there is no intra-cluster edge and all inter-cluster pairs of vertices are connected with edges of weight *max*(*w_IG_*). A *wMQ* of 1 corresponds to the case where no inter-cluster vertices are connected and all pairs of intra-cluster vertices are connected with edges of weight *max*(*w_IG_*). A high-standard consensus clustering should maximise this index.

ClustOmics computes the *wMQ* for the LCD and MCL consensus clusterings obtained with various numbers of supports thresholds and having passed the filtering steps, and returns the one that maximises this quality measure.

### Datasets and tools used for computing input clusterings

We used ClustOmics to predict cancer subtypes from gene expression, micro-RNA expression and DNA methylation datasets available in *The Cancer Genome Atlas* (TCGA) [42]. Our case-study is based on the same datasets as in *Rappoport and Shamir*’s review on multi-omic clustering methods [1]. The data covers ten cancer types: Leukemia (AML), Breast (BIC), Colon (COAD), Glioblastoma (GBM), Kidney (KIRC), Liver (LIHC), Lung (LUSC), Ovarian (OV), Sarcoma (SARC) and Skin (SKCM). For each cancer type, from 197 and up to 1098 patients were measured for at least one of the three omics (expression, miRNA and methylation), of which 170 up to 621 patients being measured for all three. More details on missing data per cancer type are given in Table 5.

To generate input clusterings to be fused by ClustOmics, we used five state-of-theart integrative clustering tools, summarised in Table 6: PINS [6], SNF [7], NEMO [8], rMKL [9] and MultiCCA [10]. The first four tools can be used in a single-omic context as well as in a multi-omic context, for which they were all designed. Though NEMO is the only tool that can handle partial data, for comparability purposes, this functionality was not used in the analyses we conducted. Each tool has been run with default parameters and based on authors’ recommendations. For the Single-to-Multi scenario, we also computed single-omic input clusterings with k-means clustering [19], for which the optimal number of clusters *k* was determined using the Silhouette index [24] with values of *k* ranging from 2 to 20.

### Computation of COCA consensus clusterings

To assess ClustOmics performance with respect to a state-of-the-art integration method based on consensus clustering, we applied COCA (Cluster-Of-Clusters Analysis) [5] on each integration scenario and from the same set of input clusterings as for ClustOmics. At the more so that COCA has already been applied to cancer-subtyping in a multi-omic context [43, 44].

COCA is an integrative clustering tool based on the Consensus Clustering (CC) algorithm introduced by Monti *et al.* [45]. The CC algorithm implements a resampling and co-occurrences-based strategy to assess the stability of clusters when analyzing a single dataset. By re-sampling multiple times a single dataset and applying a clustering algorithm on each perturbed dataset, and from the co-occurrences counts of samples in clusters, a consensus matrix is computed and used as a similarity matrix to compute a final consensus clustering. COCA was run using default parameters, under the same integration scenarios as in ClustOmics, and using the same set of input clusterings.

### Clustering pairwise similarity metric

To evaluate the similarity of ClustOmics and COCA consensus clusterings with respect to their inputs or each other, we used the Adjusted Rand Index (ARI) [20], a measure of similarity between two data clusterings. While the ARI has been used to evaluate the quality of classifications compared to ground-truth data, here we use it to compare the similarity of various clusterings, without considering any quality aspect.

### Biological metrics

To explore the biological relevance of input clusterings and consensus clustering results, we computed the *overall survival rate* of patients. As cancer acuteness is proved to be related to its molecular subtype [46, 47, 48], we further investigated whether it was significantly different across clusters using the exact log-rank test for more than two groups, introduced in [49]. For each clustering, the p-value of the log-rank test was computed using 100, 000 random permutations of the data.

In addition, we performed an analysis of *clinical labels enrichment* in clusters, using 32 labels available from TCGA metadata (see Table 7). The idea is that patients affected by the same cancer subtype should also share, to a certain extent, the same clinical characteristics. The abundance of clinical labels in clusters and their statistical over-representation provide information on the biological robustness of clusterings. To perform this analysis, we used pan-cancer (e.g. age at diagnosis or pathologic stage of cancer) and cancer-specific clinical labels for each cancer type (e.g. presence of colon polyps for colon cancer, or smoking history for lung cancer). Clinical labels that were absent for more than half of the patients were removed from the analysis. We used the *χ*^2^ test for independence for discrete parameters and the Kruskal-Wallis test for numeric parameters to assess the enrichment of the clinical labels in a cluster. To increase the robustness of the results, we applied a bootstraping strategy, computing the test on randomly permuted data to derive an empirical p-value (100, 000 permutations).

**Table 7.** Pan-cancer and cancer-specific labels used for clinical label enrichment analysis. Pathological M, N and T labels refer to the TNM staging system, which describes the anatomical extent of tumor cancers [25].

|  |  |
| --- | --- |
| Pan-cancer | Age at initial pathologic diagnosis, Gender, Pathologic M, Pathologic N, Pathologic T, Pathologic stage, Histological type, New neoplasm event type, Neoplasm histologic grade |
| AML | CALGB cytogenetics risk category, FAB morphology code |
| BIC | PAM50Call RNAseq, Estrogen receptor status, Progesterone receptor status, ER level cell percentage category, PR level cell percentage category |
| COAD | Presence of colon polyps, History of colon polyps |
| GBM | Prior glioma |
| KIRC | Hemoglobin result, Platelet qualitative result, Serum calcium result, White cell count result |
| LIHC | Adjacent hepatic tissue inflammation extent type, Albumin result specified value, Creatinine value, Fetoprotein outcome value, Fibrosis ishak score |
| LUSC | Tobacco smoking history, Number pack years smoked |
| OV | No supplementary clinical label |
| SARC | No supplementary clinical label |
| SKCM | Melanoma Clark level value, Melanoma ulceration indicator |

One has to keep in mind that molecular data does not always explain survival or clinical differences between groups of samples. Therefore, in the discussion of the results we consider survival and clinical analysis as ways to interpret patterns captured by the various clustering results, and do not favor one metric over the other.

Finally, to evaluate differentially expressed genes across consensus clusters generated using StoM scenario on BIC study case, we applied the Kruskal-Wallis test on each gene available from the BIC expression dataset. P-values were adjusted to control the False Discovery Rate (FDR) [50], filtered with a 0.001 significance threshold, and top 1000 most significant genes were kept for further analysis. Using Hierarchical Clustering [51], we clustered the top gene list and investigated clusters for enriched Gene Ontology [52] Biological Process terms with Cluster Profiler [53]. FDR-adjusted p-values were filtered with a 0.05 cutoff.

### Implementation

ClustOmics is implemented based on the Neo4j graph database management system and uses APOC and Graph Data Science Neo4j libraries. Queries on the graph atabase are performed in Cypher, Neo4j’s graph query language, and are encapsulated in Python scripts. To facilitate its use, ClustOmics can be run through the Snakemake workflow management system.

ClustOmics was tested on a desktop-computer with an Intel Xeon processor (2.70GHz, 62GB of RAM) running on Ubuntu 18.04. For the TCGA real datasets it was applied to, ClustOmics runtimes range from a few minutes for small datasets (AML, Multi-to-Multi scenario) up to 2 hours for the largest dataset (BIC, Single-to-Multi scenario), most of the computation time being consumed for the construction of the integration graph. The graph being stored in a Neo4j database, this step is only to be performed once for each integration scenario and parameters for graph filtering can be further set and tuned without re-computing the graph.

ClustOmics source code, released under MIT licence, as well as the results obtained on the ten cancer types with the two integration scenarios described in this paper are available on Github: https://github.com/galadrielbriere/ClustOmics.

## Abbreviations

AML: Acute Myeloid Leukemia
ARI: Adjusted Rand Index
BIC: Breast Invasive Carcinoma
CALGB: Cancer and Leukemia Group B
COAD: Colon Adenocarcinoma
EAC: Evidence Accumulation Clustering
ER: Estrogen Receptor
FAB: French-American-British (morphology code)
FDR: False Discovery Rate
GBM: Glioblastoma
KIRC: Kidney Renal Clear Cell Carcinoma
LCD: Louvain Community Detection
LIHC: Liver Hepatocellular Carcinoma
LUSC: Lung Squamous Cell Carcinoma
MCL: Markov Clustering
MtoM: Multi-to-Multi (integration scenario)
MQ: Modularization Quality
OV: Ovarian Carcinoma
PR: Progesterone Receptor
SARC: Sarcoma
SKCM: Skin Cutaneous Melanoma
StoM: Single-to-Multi (integration scenario)
TCGA: The Cancer Genome Atlas
wMQ: weighted Modularization Quality

## Declarations

**Ethics approval and consent to participate**

No ethics approval was required for the study.

## Consent for publication

Not applicable.

## Availability of data and materials

The datasets analysed in the study were obtained from Ron Shamir’s lab and are available at

http://acgt.cs.tau.ac.il/multi_omic_benchmark/download.html.

ClustOmics’ source code, released under MIT licence, as well as the results obtained on TCGA cancer data are available on Github: https://github.com/galadrielbriere/ClustOmics.

## Author’s contributions

GB, PT and RU conceived the project and the method on which ClustOmics is based. GB developed the ClustOmics tool and performed the bioinformatics experiments. GB, ED, PT and RU participated at the analysis of the results. GB, ED, PT and RU wrote, read and approved the final manuscript.

## Acknowledgments

We kindly thank Nora Speicher for sending us the executable of the rMKL tool.

