## Supplementary File for "Consensus clustering applied to multi-omic disease subtyping"

### Supplementary material

**Supplementary Figure 1 Distribution of clinical labels found enriched upon all cancer types : (A) in the consensus clusterings for both MtoM and StoM All scenarios, (B) in MtoM consensus clusterings and the corresponding inputs, (C) in StoM All consensus clusterings and the corresponding inputs.**

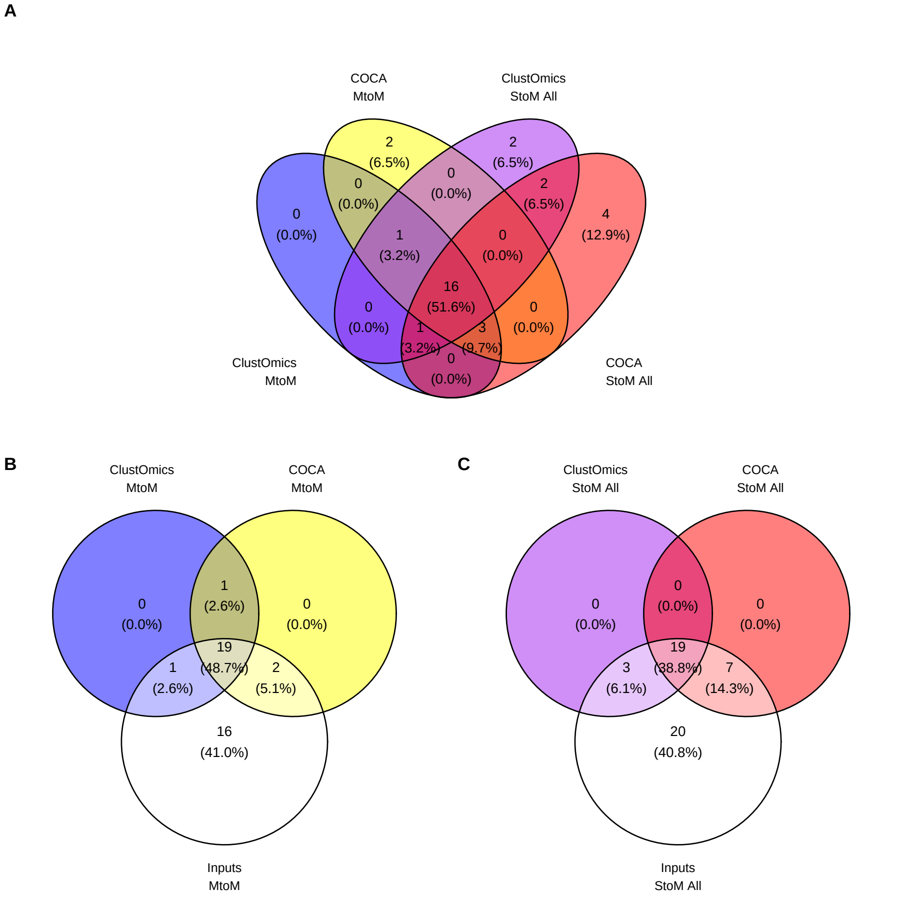

**Supplementary Figure 2 ARI heatmaps revealing input and consensus clustering similarities for the MtoM scenario, upon the ten cancer types.**

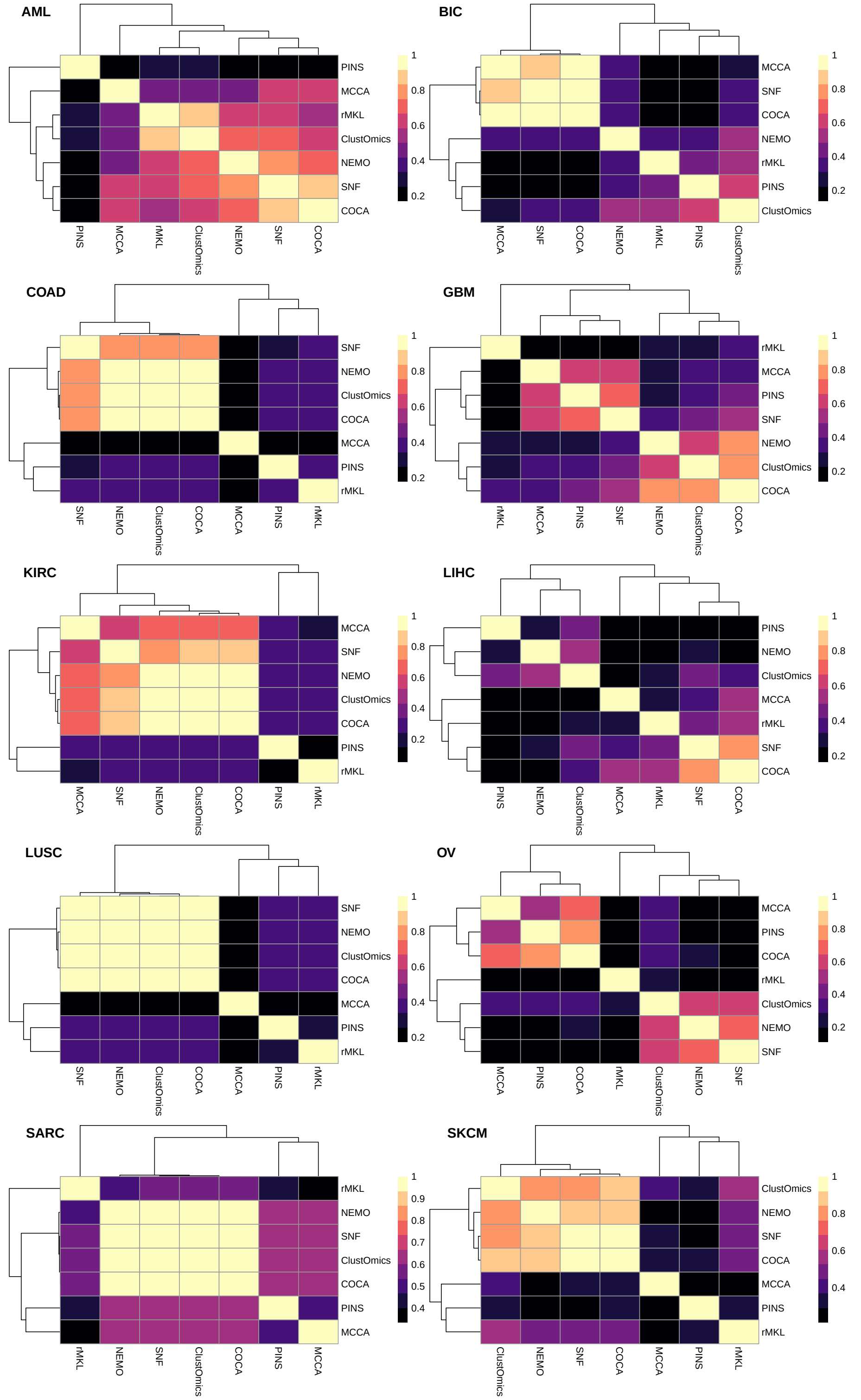

**Supplementary Figure 3 Expression heatmap of top 1000 differentially expressed genes across breast-cancer ClustOmics StoM consensus clusters.** Genes are displayed in rows and clustered according to their expression profile. Patients are displayed in columns and are ordered according to their consensus cluster attribution. PAM50 labels are given as supplementary annotations for patients.

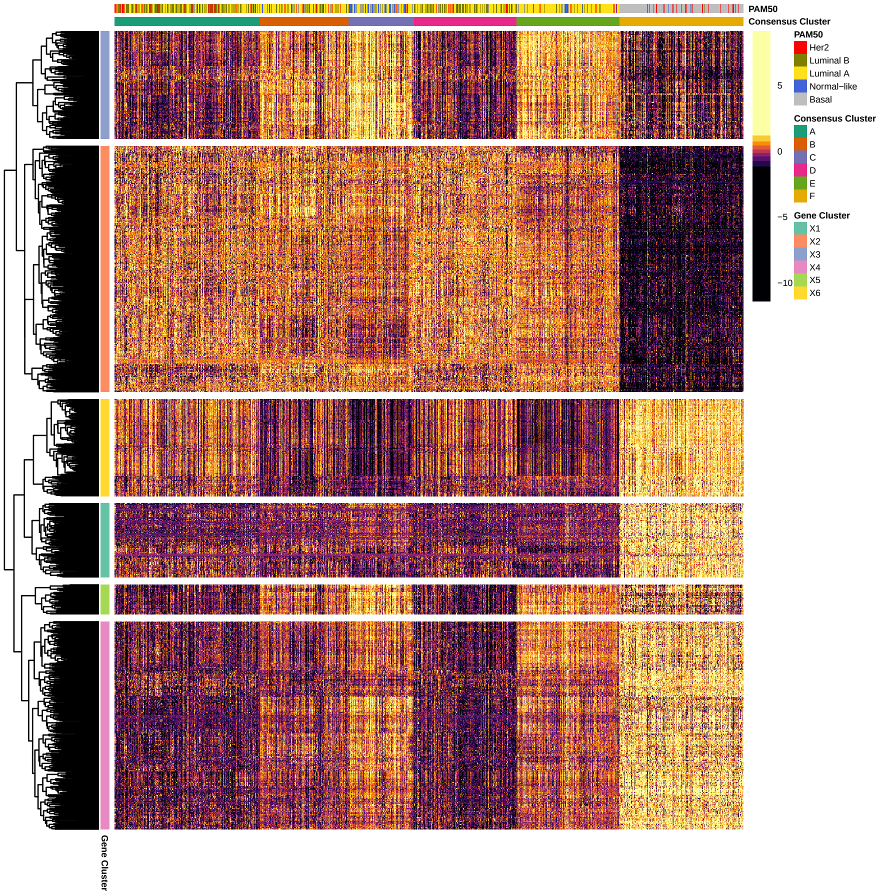

**Supplementary Figure 4 GO (Biological Process) enrichment analysis of 5 gene clusters found differentially expressed across breast-cancer StoM consensus clusters.** P-values have been adjusted using the Benjamini-Hochberg correction and filtered with a $0.05$ cutoff.

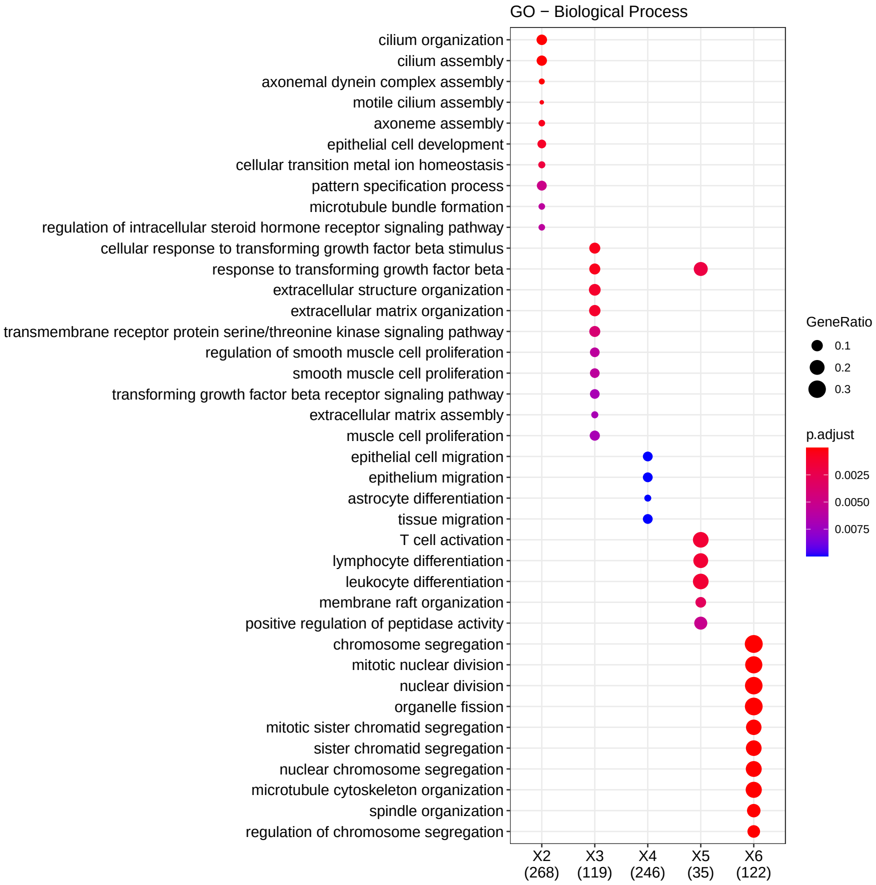
